# Bio-CM^2^: Distributed computational optics for cortex-wide cellular imaging

**DOI:** 10.64898/2026.07.27.740823

**Authors:** Guorong Hu, Qilin Deng, Tianrui Qi, Zhixiong Chen, Bradley C. Rauscher, Nathan Chai, Daria Bogatova, Bethany Weinberg, Jesse Smith, Ian G. Davison, Martin Thunemann, Anna Devor, Lei Tian

## Abstract

Understanding distributed biological systems, particularly neural circuits, requires simultaneous cellular-resolution imaging across millimeter-scale fields of view (FOV). Existing miniature microscopes remain fundamentally constrained by trade-offs among FOV, spatial resolution, and optical complexity, limiting their ability to bridge cellular microscopy with cortex-scale imaging. Here we introduce distributed computational optics, a framework that distributes image formation across coordinated optical modules and computationally integrates their measurements into a unified image. We realize this framework in Bio-CM^2^, a computational miniature mesoscope that partitions the imaging field across four optical modules while converging their measurements onto a common image sensor. This architecture overcomes the aberration-scaling limitations of conventional miniature optics while avoiding the hardware complexity of multi-camera systems and the contrast degradation associated with optical multiplexing. Bio-CM^2^ achieves a 7.5 *×* 10 mm^2^ FOV while enabling cellular-resolution *in vivo* imaging at video rates. We demonstrate its utility through two complementary imaging modalities in head-fixed mice: cortex-wide functional vascular imaging, enabling simultaneous quantification of pial arteriole vasomotion and mesoscale hemodynamic functional connectivity, and cellular-resolution calcium imaging, resolving the activity of over 3,000 neurons together with mesoscale neuronal functional connectivity. We further demonstrate the versatility of the platform through cellular-resolution imaging of entire coronal mouse brain sections, population-scale imaging of freely behaving *Caenorhabditis elegans*, and odor-evoked calcium imaging of the main olfactory bulb in head-fixed mice, highlighting its broad applicability across diverse biological systems and imaging modalities. By overcoming the conventional trade-off between FOV and spatial resolution in a compact miniature platform, Bio-CM^2^ establishes distributed computational optics as a scalable framework for multiscale biological imaging.

## 1 Introduction

Many biological processes are inherently multiscale, spanning cellular, tissue, and organ levels. Imaging these processes therefore requires simultaneous cellular resolution and large fields of view (FOVs). This challenge is particularly important in systems neuroscience, where understanding how distributed neural circuits give rise to perception, cognition, and behavior requires monitoring thousands of neurons across multiple cortical regions [1]. Although modern benchtop mesoscopes have enabled cellular-resolution imaging over large FOVs [2–6], their size and complexity limit applications requiring compact or miniature imaging systems [7]. Bridging cellular-resolution microscopy with mesoscopic imaging in a miniature platform therefore remains a longstanding challenge.

Existing miniature microscopes have largely pursued three strategies for scaling cellular-resolution imaging to larger FOVs, each with important limitations. First, conventional miniature microscopes rely on a single objective lens to image the entire FOV. As the FOV increases, off-axis aberrations rapidly degrade image quality toward the image periphery [8], requiring increasingly sophisticated optical correction that inevitably increases optical complexity, size, weight, and manufacturing difficulty. Consequently, first-generation GRIN-lens miniscopes provide excellent cellular resolution but are typically limited to FOVs smaller than 1 mm^2^ [7]. More recent miniature objectives based on compound lenses or diffractive optics extend the imaging coverage to approximately 4 × 4 mm^2^, but at the cost of substantially increased optical or fabrication complexity [9–12]. Second, camera-array architectures distribute imaging across multiple optical paths and have recently demonstrated remarkable cortex-scale recordings [13, 14]. However, replicating cameras and optical modules substantially increases packaging complexity, synchronization requirements, power consumption, and system cost. Third, computational imaging methods combine optical encoding with computational reconstruction to extend the capabilities of miniature imaging systems. Existing systems, including lensless imagers [15, 16], light-field microscopes [17, 18], and microlens-array-based architectures [19–22], such as our previous computational miniature mesoscope (CM^2^) [19–21], largely rely on optical multiplexing, whereby signals from multiple object locations are superimposed onto individual sensor pixels. While this strategy enables highly compact implementations, it inevitably mixes in-focus signals with out-of-focus background fluorescence, reducing image contrast and signal-to-noise ratio (SNR) under *in vivo* imaging conditions and fundamentally limiting measurement fidelity.

To overcome these limitations, we propose a distributed computational optics framework in which optical image formation is distributed across multiple coordinated optical modules and computationally integrated into a unified image, rather than relying on a single objective lens. Here we introduce Bio-CM^2^, a next-generation computational miniature mesoscope that implements this framework using four coordinated imaging modules whose complementary measurements are acquired simultaneously on a common image sensor without optical multiplexing (Fig. 1a,b). By distributing optical complexity across individual imaging modules, Bio-CM^2^ scales optical design with each local subfield rather than the total FOV, while computational reconstruction integrates their measurements into a seamless high-resolution image. This architecture overcomes the conventional trade-off between FOV and spatial resolution while preserving image contrast and the compact form factor required for miniature *in vivo* imaging. The resulting platform enables cellular-resolution *in vivo* imaging over a 7.5 × 10 mm^2^ FOV at video rates.

**Figure 1.**
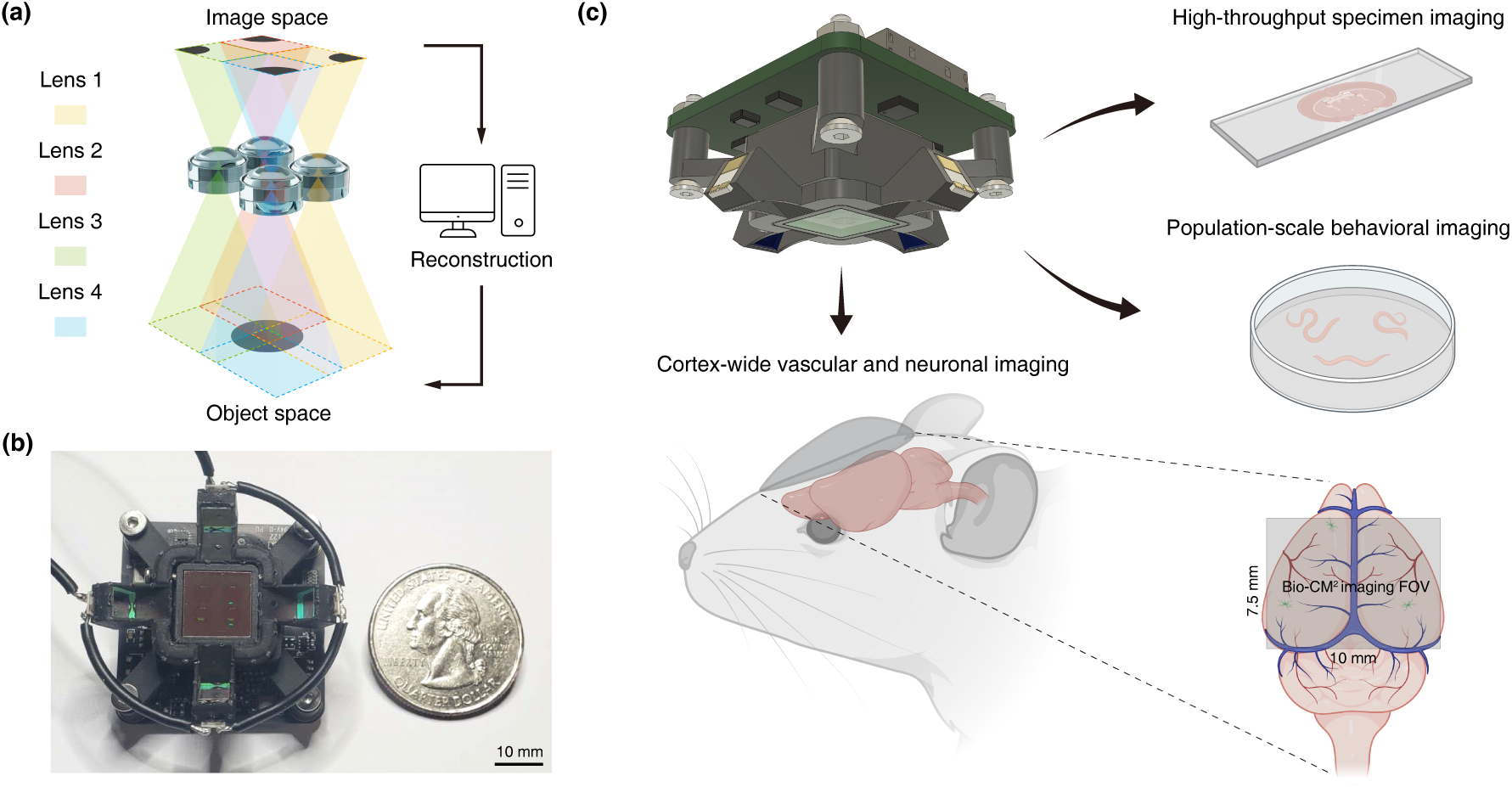
Bio-CM^2^: a miniature distributed computational imaging platform for mesoscale biological imaging. (a) Bio-CM^2^ imaging principle. A distributed array of four miniature lenses projects partially overlapping sub-images onto a common image sensor, enabling high-resolution imaging over a wide FOV through computational fusion. (b) Photograph of the Bio-CM^2^ prototype. A U.S. quarter is shown for size comparison. (c) Representative applications of Bio-CM^2^. The compact imaging platform enables high-throughput imaging of large biological specimens, population-level behavioral studies of small organisms, and cortex-wide imaging of neuronal and vascular dynamics in mice.

We first establish the distributed computational optics principles underlying the Bio-CM^2^ architecture, followed by the optical implementation, computational reconstruction framework, and quantitative characterization of its spatial resolution, FOV, and depth of field (DOF). We then evaluate Bio-CM^2^ across representative biological applications spanning the three major application areas illustrated in Fig. 1c. Specifically, we demonstrate cellular-resolution imaging of entire coronal mouse brain sections for high-throughput imaging of large biological specimens, population-scale imaging of freely behaving *Caenorhabditis elegans* for behavioral studies of small organisms, and *in vivo* imaging in head-fixed mice through quantitative functional vascular imaging spanning pial arteriole vasomotion and mesoscale hemodynamic functional connectivity, cortex-wide calcium imaging resolving over 3,000 neurons together with mesoscale neuronal functional connectivity, and odor-evoked calcium imaging in the main olfactory bulb. Together, these results establish Bio-CM^2^ as a platform for multiscale biological imaging and demonstrate the potential of distributed computational optics as a scalable strategy for extending cellular-resolution imaging to mesoscale FOVs within compact miniature imaging systems.

## 2 Results

### 2.1 Principle of Bio-CM**^2^**

Bio-CM^2^ adopts a distributed optical architecture to bridge cellular-resolution microscopy with millimeter-scale imaging in a compact miniature platform. As illustrated in Fig. 1a, the imaging field is partitioned into four partially overlapping subfields, each captured by an independent miniature optical module arranged in a 2 × 2 configuration. The four views are subsequently reconstructed into a single seamless wide-FOV image through view-dependent computational correction.

Because each optical module images only a fraction of the total FOV, the maximum field angle is substantially reduced compared with a conventional single-objective design. This distributed imaging strategy alleviates off-axis aberrations while enabling a substantially larger imaging area without increasing optical complexity.

Unlike previous computational miniature microscopes, Bio-CM^2^ partitions the FOV *before* image formation rather than relying on optical multiplexing followed by computational demultiplexing. Each sensor pixel therefore receives light from only one object-space location within a single imaging channel, preserving native image contrast while avoiding the loss of measurement fidelity associated with optical multiplexing under *in vivo* imaging conditions.

Following acquisition, each imaging channel is independently calibrated to compensate for geometric distortion, illumination nonuniformity, and spatially varying blur. The corrected views are then registered and stitched into a seamless wide-FOV image with uniform image quality across the entire imaging area (Fig. 2).

**Figure 2.**
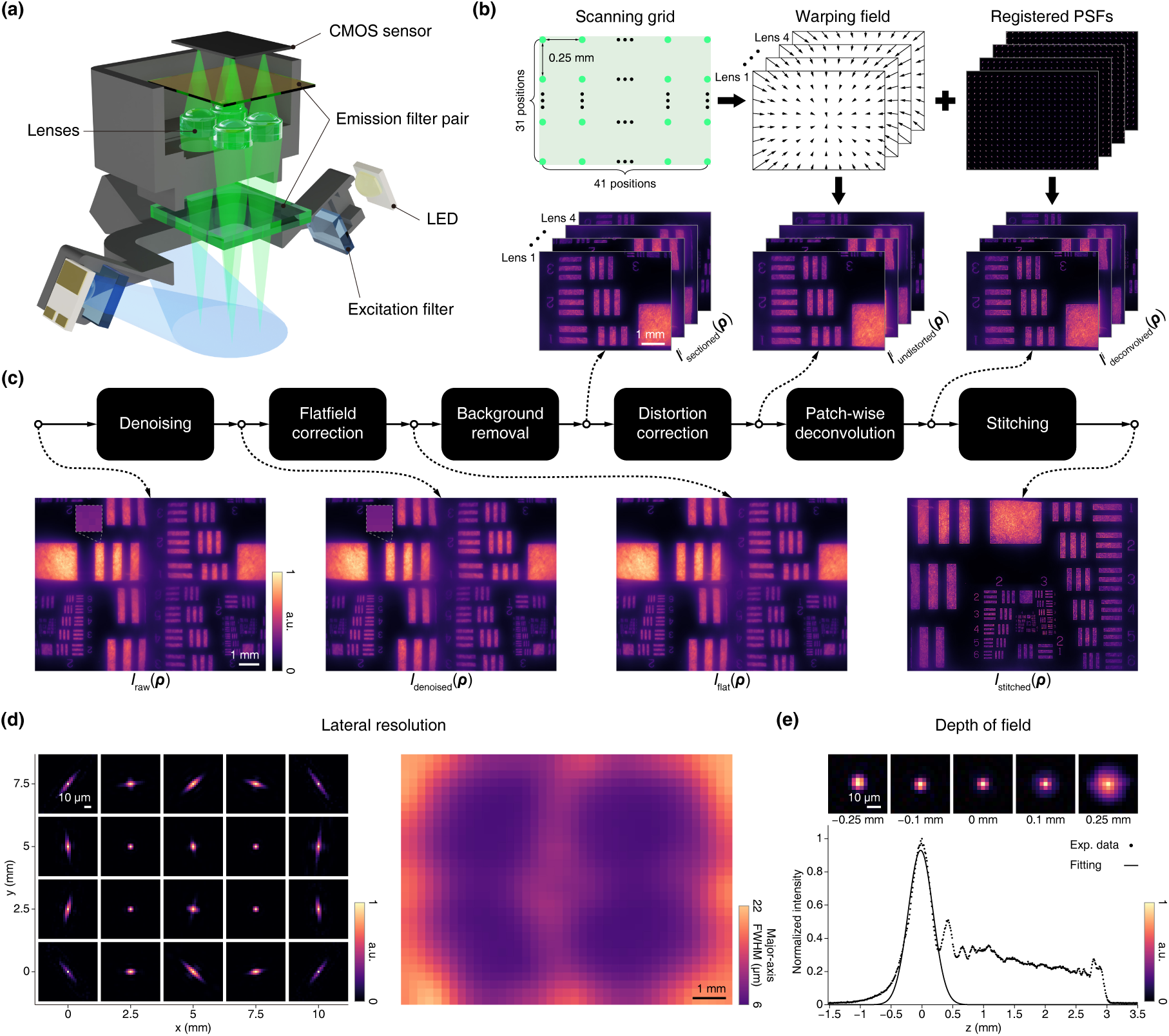
Bio-CM^2^ system architecture, computational reconstruction, and imaging characterization. a) Exploded view of Bio-CM^2^. The system consists of a four-lens imaging module and a four-LED illumination module. (b) PSF calibration and geometric registration. Experimentally measured PSFs acquired over a calibration grid are used to estimate lens-specific warping fields and generate registered PSF libraries for image reconstruction. (c) Computational reconstruction pipeline. Raw measurements are sequentially processed by denoising, flat-field correction, optional background removal, geometric distortion correction, patch-wise deconvolution, and image stitching to produce the final reconstructed image. (d) Lateral resolution characterization. Left, representative PSFs sampled across the imaging FOV. Right, spatial map of the conservatively reported major-axis FWHM obtained by rotated anisotropic Gaussian fitting. (e) Axial response characterization. Top, representative on-axis PSFs measured at different depths. Bottom, measured axial intensity profile with the Gaussian fit used to estimate the DOF.

### 2.2 Implementation and characterization of Bio-CM**^2^**

The assembled Bio-CM^2^ device is shown in Fig. 2a. The system integrates a 2 × 2 array of miniature imaging lenses together with four symmetrically positioned LEDs that surround the lens array. The four imaging channels cooperatively acquire complementary subfields that are computationally reconstructed into a seamless 7.5 mm × 10 mm FOV with cellular-scale spatial resolution. The illumination module provides near-uniform fluorescence excitation across the entire imaging region, which is critical for reliable *in vivo* imaging. The emitted fluorescence is collected through a shared emission filter assembly onto a CMOS sensor.

Following assembly, the imaging module is experimentally calibrated using a benchtop characterization system. Spatially varying point spread functions (PSFs) are measured by scanning a fluorescent bead over a grid (Fig. 2b and Supplementary Section 1.4). The expected PSF positions are first predicted from a geometric optics model and then compared with the experimentally measured PSF locations. The resulting spatial displacement field defines the lens-specific geometric distortion map. Together, the calibrated PSFs and distortion maps constitute the system model for subsequent image reconstruction.

Starting from the raw measurements, Bio-CM^2^ reconstructs images through the computational pipeline illustrated in Fig. 2c. The raw images are first denoised and corrected for illumination nonuniformity, followed by optional background subtraction, geometric distortion correction, patch-wise deconvolution using experimentally measured PSFs, and image stitching. Representative intermediate results illustrate the contribution of each processing stage, with the inset highlighting denoising performance for a representative 4 × 4 pixel image patch.

We next characterized the lateral resolution using the experimentally calibrated PSFs (Fig. 2d). For object-space regions imaged by multiple lenses, the smallest PSF was selected to represent the local resolution. Representative PSFs sampled across the imaging FOV and the corresponding spatial map of the (worst-case) major-axis full-width at half-maximum (FWHM), obtained by fitting a rotated anisotropic Gaussian model, are shown in Fig. 2d. Near the optical axis of each imaging module, the PSFs are nearly isotropic with a FWHM of approximately 6 µm. At larger field angles, off-axis aberrations introduce increasing anisotropy, and the conservatively reported major-axis FWHM gradually increases to approximately 22 µm near the lens periphery. Given an object-space sampling of 2.89 µm per pixel and a nominal numerical aperture of 0.17, image quality is sampling-limited near the lens centers and becomes aberration-limited toward the edge of each imaging module.

We further characterized the axial response by translating a fluorescent bead over a 5-mm axial range in 10 µm increments (Fig. 2e). Representative PSFs at different depths and the corresponding axial intensity profile are shown in the upper panel. Owing to spherical aberration, the axial response is asymmetric, exhibiting an extended tail when defocusing away from the lens. Gaussian fitting of the main lobe yields an estimated DOF of approximately 473 µm.

The FOV was quantified using a fluorescent resolution target masked to different effective imaging areas. The four imaging channels remain completely separated on the sensor for object sizes up to the designed 7.5 × 10 mm^2^ FOV. As the object size exceeds this limit, adjacent sub-images begin to overlap, thereby defining the maximum non-overlapping FOV of the system.

Finally, we compared Bio-CM^2^ with representative miniature imaging systems [4, 9–12, 14–16, 19, 20, 22, 23] by plotting FOV area against the inverse of the reported best lateral resolution (Extended Data Fig. 1, Supplementary Section 1.8). The background contours denote a FOV–resolution figure of merit, defined as the FOV area divided by the square of the best reported lateral resolution, and are included for visual comparison only. Bio-CM^2^ combines a large FOV with high spatial resolution, placing it near the upper-left region of the comparison space.

### 2.3 High-throughput imaging of fixed brain sections

To demonstrate the capability of Bio-CM^2^ for high-throughput imaging of large biological specimens, we imaged an entire 75-µm-thick fixed coronal mouse brain section with GFP expression targeted to the medial amygdala (MEA) (Fig. 3; raw measurement shown in Supplementary Fig. 14(a)). The reconstructed image spans the complete coronal section in a single acquisition, encompassing multiple major brain regions, including the neocortex (CTX), hippocampal formation (HPF), thalamus (TH), hypothalamus (HYP), piriform cortex (PIR), basolateral amygdala (BLA), and medial amygdala (MEA). The full reconstruction is displayed using a square-root intensity transform to simultaneously visualize low-intensity tissue background and bright neuronal structures, whereas representative enlarged regions are shown with linear intensity scaling.

**Figure 3.**
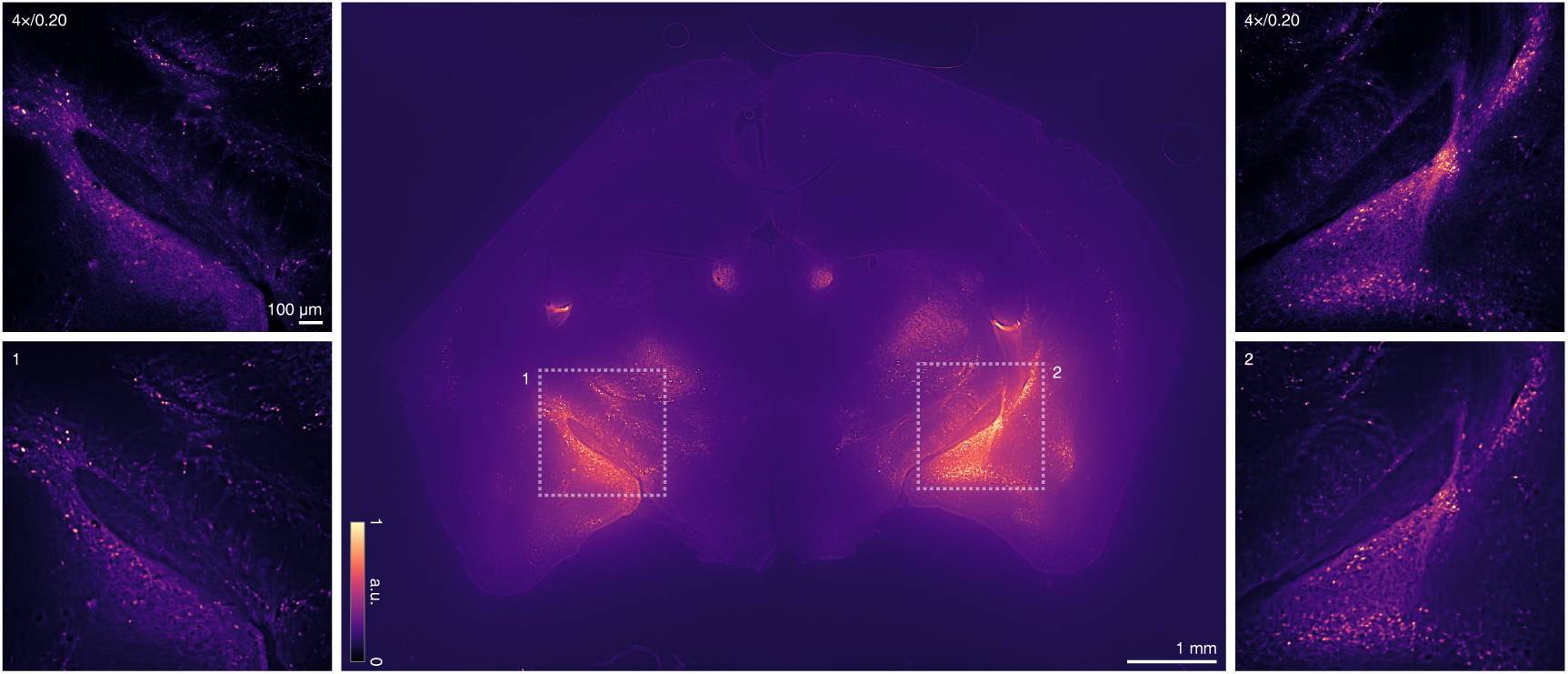
Single-shot imaging of a centimeter-scale mouse brain section with Bio-CM^2^. Middle, Bio-CM^2^ reconstruction of an entire 75-µm-thick fixed coronal mouse brain section spanning multiple major brain regions, acquired in a single measurement. Bottom left and right, representative magnified views highlighting neuronal morphology. Top left and right, corresponding reference images acquired using a conventional 4*×*/0.20 NA objective.

We assessed reconstruction fidelity by comparing representative regions of interest (ROIs) with corresponding images acquired using a conventional 4×/0.20 NA objective (top left and top right). Bio-CM^2^ accurately recovers neuronal morphology and fluorescence intensity patterns that closely match the reference measurements.

These results demonstrate that Bio-CM^2^ enables scan-free cellular-resolution imaging across entire coronal mouse brain sections, combining tissue-scale coverage with sufficient spatial resolution for neuronal morphology. This capability enables rapid screening and quantitative analysis of large histological specimens spanning multiple anatomical regions within a single acquisition.

### 2.4 Population-scale imaging of *C. elegans*

Bio-CM^2^ is also well suited for population-scale imaging of small organisms. To demonstrate this capability, we imaged freely behaving *C. elegans* expressing YFP-tagged expanded polyglutamine (polyQ) protein in body-wall muscle cells, a widely used model for studying protein aggregation associated with Huntington’s disease. The reconstruction is shown in the center panel of Fig. 4 (raw measurement shown in Supplementary Fig. 14(b)), with four representative enlarged views displayed at the corners. Operating at video rates over a centimeter-scale FOV, Bio-CM^2^ simultaneously captures numerous independently behaving worms exhibiting diverse body postures while resolving individual fluorescent polyQ aggregates within each animal. This unique combination of large-area coverage and cellular-resolution imaging enables high-content phenotypic analysis by directly correlating locomotor behavior with intracellular protein aggregation across large populations under identical experimental conditions, without the need for immobilization.

**Figure 4.**
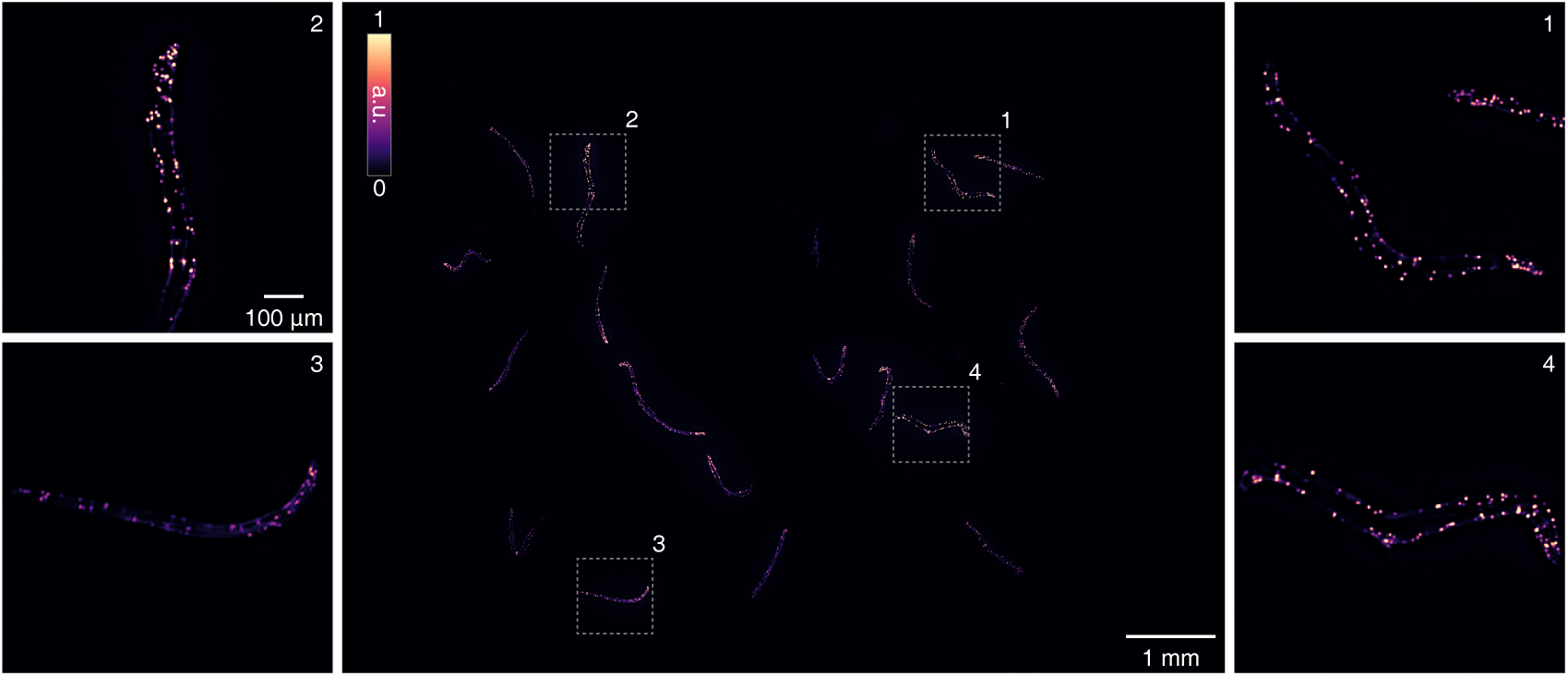
Bio-CM^2^ enables simultaneous observation of multiple freely behaving small model organisms. A population of fluorescently labeled *C. elegans* is imaged in a single acquisition. The enlarged side panels show representative worms from different locations across the FOV, revealing distinct body postures and subcellular polyQ-YFP protein aggregates in body-wall muscle cells.

### 2.5 Cortex-wide vascular imaging from single-vessel dynamics to functional connectivity

To demonstrate the versatility of Bio-CM^2^ for *in vivo* functional imaging, we first performed cortex-wide fluorescence imaging of the cortical surface vasculature through a bilateral chronic glass cranial window replacing the dorsal cranium, following a previously described procedure [24] (Extended Data Fig. 3a). Following intravenous injection of a dextran-conjugated fluorescent vascular dye, Bio-CM^2^ captured vascular dynamics across the entire dorsal cortex. Imaging such a large cortical area presents an additional challenge because the curved cortical surface introduces parallax between adjacent Bio-CM^2^ views. To compensate for this effect, overlapping sub-images were aligned using image warping based on manually selected corresponding feature points before stitching (Supplementary Section 2.2), enabling seamless reconstruction of the full cortical surface vasculature (Fig. 5a). The reconstructed video provides a single measurement from which vascular dynamics can be analyzed across multiple spatial scales (a representative raw measurement shown in Supplementary Fig. 14(c)).

**Figure 5.**
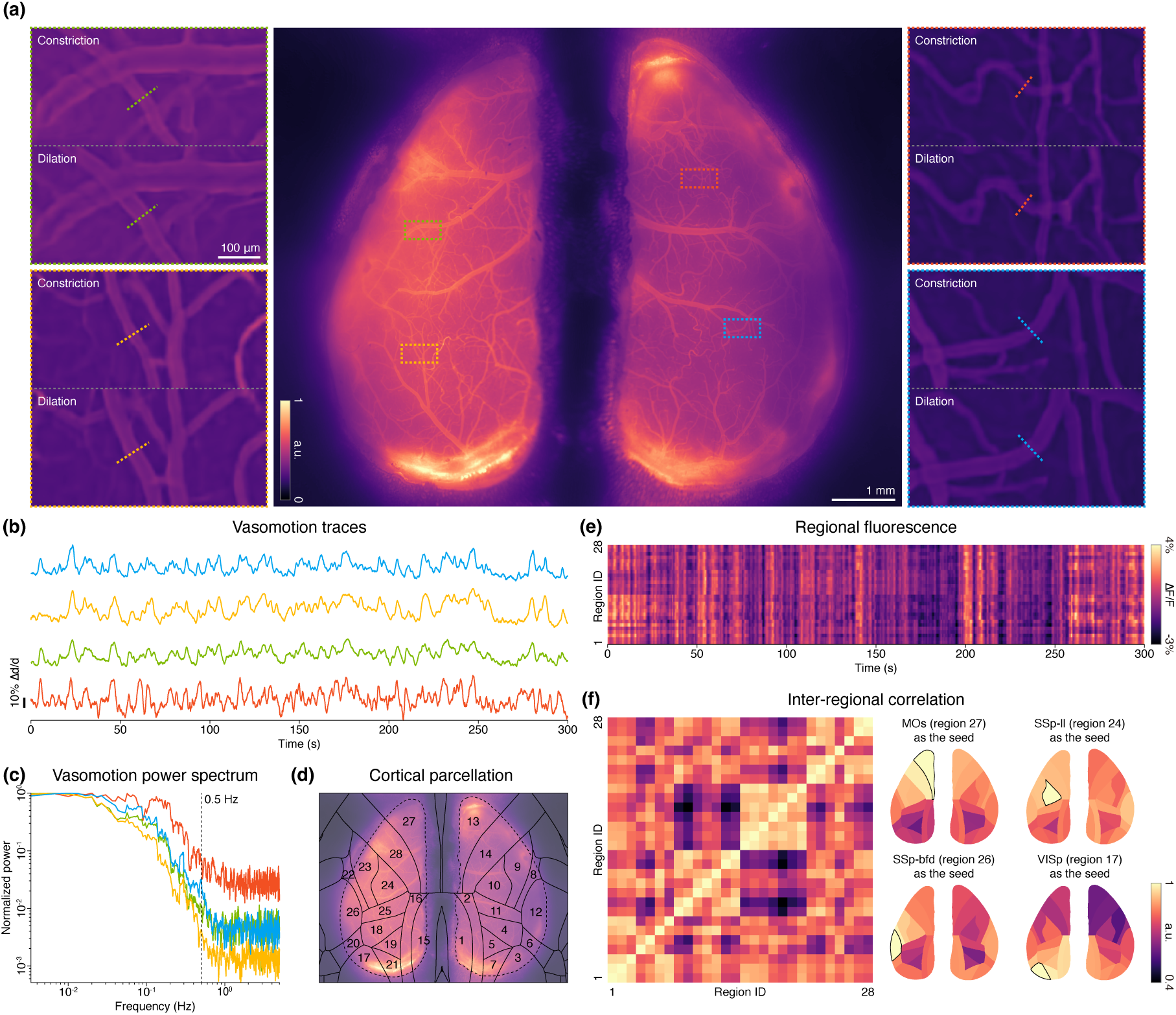
Bio-CM^2^ enables multiscale vascular imaging in head-fixed mice from single-vessel dynamics to cortex-wide hemodynamic networks. (a) Representative cortex-wide vascular image acquired through a chronic cranial window. Insets show enlarged views of representative vessel segments exhibiting vasoconstriction and vasodilation over time. The full-field image is displayed without background removal, whereas the insets are shown after background removal to enhance vessel-edge visualization. (b) Time-series traces of the relative vessel diameter changes (Δ*d/d*) measured from the representative vessel segments shown in the insets of (a). (c) Normalized power spectra of the vasomotion signals shown in (b). (d) Anatomical parcellation of the dorsal cortex based on the Allen Mouse Brain Atlas. The dashed line outlines the manually selected imaged brain region; region-wise fluorescence signals were extracted only from pixels within each cortical parcel and the outlined brain region. (e) Regional hemodynamic activity (Δ*F/F* ) quantified from the anatomically defined cortical parcels shown in (d) over the imaging session. (f) Cortex-wide functional connectivity of hemodynamic activity. Left, pairwise correlation matrix computed from the regional activity in (e). Right, representative seed-based functional connectivity maps.

At the single-vessel level, changes in pial arteriole diameter provide a direct readout of local vascular dynamics. Bio-CM^2^ resolves individual pial arterioles across the dorsal cortex, enabling quantitative measurement of these diameter fluctuations [25]. For this analysis, background removal was applied only to enhance vessel contrast for accurate vessel-edge detection during diameter quantification (the four enlarged views in Fig. 5a and Fig. 5b,c). Representative vessel segments from four cortical regions are shown in the enlarged views of Fig. 5a, where both vasoconstriction and vasodilation are observed during the recording. Vessel diameters were extracted from manually selected ROIs, and the resulting time series of relative diameter changes (Δ*d/d*) are shown in Fig. 5b. Their corresponding power spectra (Fig. 5c) reveal vasomotion frequencies predominantly below 0.5 Hz, consistent with previous reports [26, 27]. Additional vessel measurements are presented in Supplementary Section 3. Together, these results demonstrate simultaneous quantification of single-vessel dynamics spanning the entire dorsal cortex within a single recording.

At the mesoscale, the same reconstructed video was analyzed without background removal. In contrast to vessel-diameter analysis, preserving the diffuse intravascular FITC-dextran fluorescence retains low-frequency intensity fluctuations arising from the depth-integrated, optically weighted vascular signal. Because the measured fluorescence represents a depth-integrated, optically weighted signal arising from labeled blood throughout the imaging volume, its temporal fluctuations provide a readout of regional cerebral blood-volume dynamics [28]. The resulting regional fluorescence time series capture mesoscale hemodynamic activity and enable functional connectivity analysis [24]. Using the Allen Mouse Brain Atlas registered to the imaging field (Fig. 5d), we extracted regional Δ*F/F* signals from anatomically defined cortical parcels within the manually delineated brain region imaged by Bio-CM^2^ (Fig.5e), providing a mesoscale representation of hemodynamic activity across the dorsal cortex. Pairwise correlation analysis of these regional signals revealed cortex-wide hemodynamic functional connectivity (Fig. 5f). Representative seed maps derived from the MOs, SSp-ll, SSp-bfd, and VISp parcels exhibit strong bilateral symmetry and progressively decreasing correlations with increasing anatomical distance, consistent with previous mesoscale imaging studies [24, 29]. Together, these results demonstrate that Bio-CM^2^ enables simultaneous multiscale vascular imaging, bridging quantitative single-vessel measurements with cortex-wide hemodynamic functional network analysis within a single acquisition.

### 2.6 Cortex-wide calcium imaging from single neurons to mesoscale cortical dynamics

We next demonstrate the capability of Bio-CM^2^ for cortex-wide functional calcium imaging in vivo. Mice expressing GCaMP6s in vasointestinal peptide (VIP)-expressing interneurons were implanted with the same bilateral chronic glass cranial window described above [24] (Extended Data Fig. 3a). Bio-CM^2^ simultaneously captures local neuronal activity together with global cortical dynamics within a single recording, enabling cellular-resolution imaging over a centimeter-scale FOV.

Full-cortex single-neuron calcium imaging presents a demanding signal-extraction problem because neuronal activity is superimposed on strong out-of-focus fluorescence, spatially structured vascular signals, and sensor noise. Unlike the static background encountered in structural imaging, the out-of-focus fluorescence varies slowly over time compared to calcium dynamics, yet its fluctuations can be comparable in magnitude to single-neuron calcium responses. Without explicitly modeling this dynamic background, these fluctuations can be misidentified as neuronal activity. This challenge is further compounded by the low-cost CMOS sensor used in Bio-CM^2^, which exhibits higher read noise than the low-noise scientific CMOS cameras commonly used in tabletop cortex-wide imaging systems [2, 3]. We therefore first denoise the raw videos using SUPPORT [30], a state-of-the-art self-supervised learning method for *in vivo* neural imaging. This step is essential for stabilizing the subsequent source-extraction procedure; without denoising, CNMF-E frequently fails to identify reliable neuronal components.

The denoised videos, reconstructed using the standard pipeline in Fig. 2 without the static background-removal step, are then analyzed using CNMF-E [31, 32]. CNMF-E jointly separates neuronal sources from residual noise and spatially and temporally varying background fluorescence. The identified candidate neuronal components are subjected to additional quality control, including restriction to manually annotated cortical regions and filtering based on spatial correlation with the raw fluorescence video. This screening is particularly important near blood vessels, where motion and hemodynamic fluctuations quantified in Section 2.5 can generate false-positive components. Detailed procedures are provided in Supplementary Section 4.1.

Figure 6a summarizes the cortex-wide calcium imaging performance of Bio-CM^2^ (a representative raw measurement shown in Supplementary Fig. 14(d)). A pixel-wise temporal standard deviation projection of the recording reveals activity across the entire dorsal cortex, while four representative enlarged views demonstrate that individual neurons are clearly resolved in both hemispheres. To estimate the effective imaging depth of Bio-CM^2^ and validate neuron detection at the single-cell level, we performed independent two-photon imaging of the corresponding cortical region. By comparing maximum-intensity projections (MIPs) generated from different depth ranges of the two-photon image stack, we found that the projection over a depth of 100–300 µm below the brain surface exhibited the closest agreement with the Bio-CM^2^ reconstruction (Supplementary Section 5), with a high degree of correspondence in the spatial distribution and morphology of individual neurons. These results indicate that the long DOF of Bio-CM^2^ enables single-shot imaging of neuronal activity across most of layer 2/3 as a depth-integrated 2D fluorescence projection, while preserving single-neuron resolution under *in vivo* scattering conditions.

**Figure 6.**
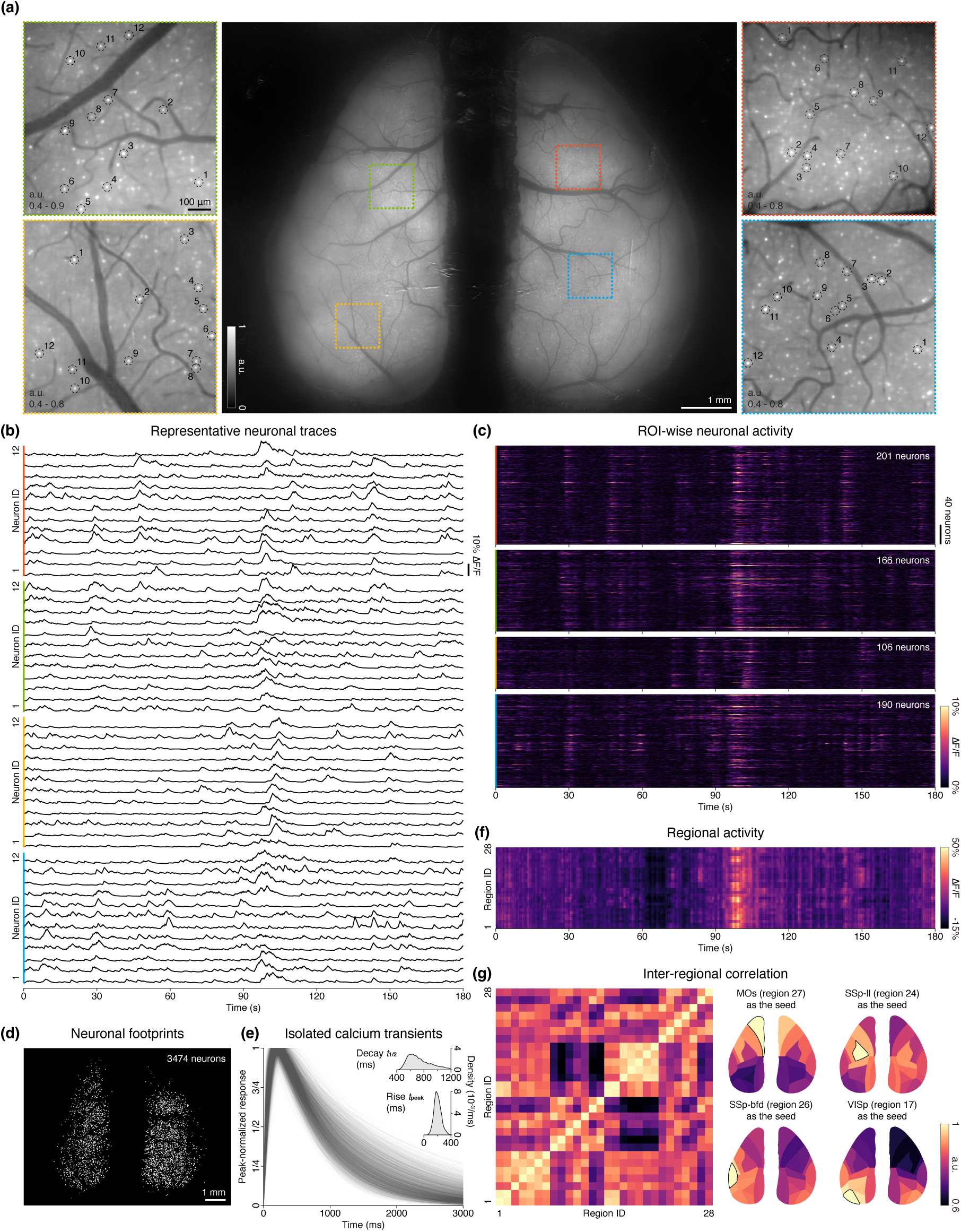
Bio-CM^2^ enables multiscale neuronal calcium imaging in head-fixed mice from single-neuron activity to cortex-wide functional networks. (a) Pixel-wise temporal standard deviation projection of the cortex-wide calcium imaging video. Insets show representative regions across both hemispheres, demonstrating that individual VIP neurons can be resolved throughout the full FOV. (b) Representative calcium traces extracted by CNMF-E from neurons in the four enlarged regions in (a). (c) Neuronal activity matrices corresponding to the four enlarged regions in (a). (d) Spatial footprints of all neurons identified by CNMF-E across the dorsal cortex, yielding a total of 3,474 neurons. (e) Individual calcium transients identified by CNMF-E, approximately aligned by their estimated onset. Inset, distributions of the extracted time-to-peak (*t*_peak_) and post-peak half-decay time (*t*_1_*_/_*_2_), demonstrating agreement with reported *in vivo* GCaMP6s kinetics. (f) Mesoscale neuronal activity quantified by spatially averaging fluorescence signals within anatomically defined cortical parcels. (g) Cortex-wide functional connectivity derived from the regional activity in (f). Left, pairwise correlation matrix between cortical parcels. Right, representative seed-based functional connectivity maps.

Following denoising, background separation, source extraction, and quality control, CNMF-E identified 3,474 neurons across the full FOV. Representative calcium traces are shown in Fig. 6b. The complete activity matrices from the four enlarged views are shown in Fig. 6c, illustrating the diverse temporal activity patterns of local neuronal populations. The spatial footprints of all identified neurons across the dorsal cortex are shown in Fig. 6d, highlighting the dense cortex-wide neuronal sampling achieved by Bio-CM^2^. Beyond identifying active neurons, the extracted fluorescence traces enable quantitative characterization of neuronal calcium dynamics across the cortex. Specifically, we quantified the time to peak, *t*_peak_, and the post-peak half-decay time, *t*_1*/*2_, of isolated calcium transients identified by CNMF-E. The resulting distributions (Fig. 6e) exhibit a median *t*_peak_ of 200.0ms (interquartile range (IQR), 168.5–236.7ms) and a median *t*_1*/*2_ of 703.8ms (IQR, 609.7–851.6ms), consistent with the reported *in vivo* fluorescence kinetics of GCaMP6s [33]. Together, these results demonstrate that Bio-CM^2^ supports quantitative, simultaneous functional recording from thousands of individual neurons across the dorsal cortex within a single acquisition.

Beyond single-neuron analysis, the same reconstructed calcium-imaging video can be analyzed at the mesoscale using the same anatomical parcellation framework as the vascular imaging analysis. Specifically, the reconstructed fluorescence video is partitioned into predefined cortical regions, and region-wise Δ*F/F* time courses are obtained by spatially averaging the fluorescence signals within each parcel. The resulting cortex-wide activity heatmap (Fig. 6f) reveals coordinated neuronal dynamics across the dorsal cortex. Pairwise correlation analysis and seed-based functional connectivity mapping (Fig. 6g) identify large-scale activity patterns that show a degree of similarity to those observed in the vascular imaging analysis (Fig. 5f). This correspondence is consistent with the well-established neurovascular coupling between neuronal activity and cerebral hemodynamics [34], together with the role of VIP interneurons in regulating cortical blood flow through vasodilation [35]. Together, these results demonstrate that Bio-CM^2^ supports seamless analysis across spatial scales, from single-neuron activity to cortex-wide functional organization, within a single recording, while providing a promising platform for investigating cortex-wide neurovascular coupling [36].

### 2.7 Odor-evoked calcium imaging in the main olfactory bulb

To demonstrate that Bio-CM^2^ is readily applicable beyond cortex-wide imaging, we next performed *in vivo* calcium imaging of the main olfactory bulb (MOB) of mice, where second-order neurons selectively express GCaMP6f (Extended Data Fig. 2 and Extended Data Fig. 3b). Owing to the smaller size of the MOB, only a single Bio-CM^2^ sub-image was required for imaging (Extended Data Fig. 2a). We then presented two odorants, pentyl acetate and benzaldehyde, separately to the mouse and observed distinct glomerular activation patterns for each stimulus (Extended Data Fig. 2d,e), consistent with the established odor-specific functional organization of the MOB [37].

To quantify these responses, we identified 92 glomerular ROIs based on the activity projection (Extended Data Fig. 2b) and extracted their Δ*F/F* time courses, summarized as activity heatmaps in Extended Data Fig. 2c. Representative glomerular traces further illustrate the distinct temporal response profiles evoked by the two odorants (Extended Data Fig. 2d,e). Together, these results demonstrate that Bio-CM^2^ can resolve stimulus-specific functional activity in anatomically distinct neural circuits, highlighting its versatility for *in vivo* calcium imaging beyond the cerebral cortex.

## 3 Summary and discussions

In summary, we developed Bio-CM^2^, a computational miniature mesoscope that distributes optical image formation across coordinated imaging modules and computationally integrates their measurements to achieve cellular resolution over a centimeter-scale FOV. Built from low-cost off-the-shelf components, Bio-CM^2^ was validated across diverse biological applications spanning entire coronal mouse brain sections, populations of freely behaving *C. elegans*, and cortex-wide *in vivo* functional vascular and neuronal imaging in mice. Together, these results establish Bio-CM^2^ as a versatile platform for multiscale biological imaging and demonstrate distributed computational optics as a scalable framework for extending cellular-resolution imaging to mesoscale FOVs, enabling quantitative interrogation of biological systems across spatial scales within compact miniature imaging platforms.

Although Bio-CM^2^ was demonstrated in head-fixed mice, this limitation is imposed by the current imaging electronics rather than the distributed optical architecture. The optical and computational framework introduced here is fully compatible with ongoing advances in lightweight sensor boards, compact camera modules, and flexible interconnects [9–11], providing a clear path toward wearable implementations without fundamental changes to the imaging architecture.

An important future direction is extending Bio-CM^2^ to multimodal imaging. Recent advances in multicolor and multimodal miniature microscopes have enabled simultaneous imaging of complementary biological signals within conventional miniature imaging architectures [11, 23, 38]. Building on these advances, integrating spectrally separated excitation, emission optics, and temporally synchronized illumination into Bio-CM^2^ would enable simultaneous imaging of multiple neuronal populations or concurrent recording of neuronal calcium activity together with vascular fluorescence or intrinsic optical signals over centimeter-scale FOVs. Although such integration represents a substantial systems-engineering effort, it would extend multicolor and multimodal miniature imaging from local circuits to cortex-scale networks, enabling investigation of interactions between distributed neuronal populations, cortex-wide neurovascular coupling, and concurrent hemodynamic measurements for quantitative correction of fluorescence signals [39].

From a computational perspective, the current reconstruction pipeline was designed to prioritize robustness and generality rather than maximize reconstruction quality. Two immediate opportunities exist for further improvement. First, the current patch-wise Tikhonov deconvolution could be replaced by recently developed deep learning approaches for shift-variant deconvolution [21, 40], which have demonstrated state-of-the-art reconstruction fidelity and computational efficiency in related problems. Second, the dynamic background removal used for *in vivo* calcium imaging could be substantially improved using deep learning methods developed for widefield one-photon imaging, which have demonstrated superior suppression of background fluorescence and enhancement of neuronal signals [41]. Because these algorithmic advances operate directly on the reconstructed measurements, they can be incorporated without modification to the optical hardware, further improving the performance of Bio-CM^2^ for large-scale functional imaging.

## Methods

### Hardware implementation

#### Bio-CM**^2^** Hardware

Bio-CM^2^ comprises illumination and imaging modules. Unlike our previous CM^2^ designs [19, 20], the imaging module adopts a distributed optical architecture consisting of a 2 × 2 array of miniature plano-aspheric lenses (3 mm diameter, 3.52 mm focal length; Edmund Optics #16-684) operating in a finite-conjugate configuration with approximately 0.64× magnification. To ensure low cost and reproducibility, the system was implemented using commercially available miniature optics. Among three candidate miniature lenses with similar aperture and focal length, the selected plano-aspheric lens exhibited the smallest off-axis PSF broadening across the designed sub-FOV when oriented with the plano surface facing the object (Supplementary Section 1.2).

The optical configuration provides a nominal working distance of approximately 9 mm, defined as the distance from the back surface of the lens to the object in-focus plane. This compact yet practical working distance is well suited for in vivo mouse imaging, accommodating the thickness of the cranial window while providing sufficient clearance to image neuronal activity located several hundred micrometers below the cortical surface.

The imaging geometry was co-optimized according to the target FOV, lens aperture, CMOS sensor size, and available miniature lens focal lengths (Supplementary Section 1.1). The four lenses project non-overlapping sub-images onto a shared backside-illuminated CMOS sensor (4000×3000 pixels, 1.85 µm pixel size; The Imaging Source DMM 37UX226-ML) without light barriers between adjacent lenses. Lens positions were optimized to maximize the imaging FOV while preventing sub-image overlap, resulting in a designed FOV of 7.5 × 10 mm^2^ (Supplementary Section 1.1).

The emission filter stack follows our previous CM^2^ V2 design [20] (Supplementary Section 1.5). Fluorescence emission passes through an interference filter (Chroma Technology ET525/50m) followed by a long-pass absorption filter (Edmund Optics #54-467) to suppress excitation leakage. The complete imaging module is assembled using custom 3D-printed holders (black resin V4.1; Formlabs Form 3+).

The illumination module builds upon our previous CM^2^ V2 design [20] while simplifying the implementation (Supplementary Section 1.6). Four identical illumination units are positioned symmetrically at an approximately 30^◦^ elevation angle around the imaging array, each comprising a surface-mounted LED (470 nm center wavelength; Lumileds LXML-PB01-0040) and an excitation filter (Chroma Technology ET470/40x). Unlike the previous design, Bio-CM^2^ relies directly on the built-in hemispherical lens of the surface-mounted LED package to shape the illumination, eliminating the need for additional illumination optics. This simplified architecture reduces system complexity while maintaining homogeneous excitation across the entire imaging FOV. The maximum irradiance at the sample plane reaches 1 mW/mm^2^ under a 1 A drive current.

A complete bill of materials, including part numbers and system cost, is provided in Supplementary Section 1.7.

#### Benchtop system

The benchtop system is used for optical alignment, PSF calibration, and performance validation. Once calibrated, Bio-CM^2^ operates as a stand-alone imaging device. The system (Supplementary Section 1.8) consists of one illumination path and two imaging paths separated by a fluorescence filter set (Thorlabs MF469-35, MF525-39, MD498).

Excitation light from a 470 nm LED (Thorlabs M470L5) is collimated (ACL12708U-A), homogenized with a diffuser (Edmund Optics #47994), and relayed to the objective pupil using a 4*f* system (Thorlabs LA1986-A and LA1433-A). Samples are mounted on a six-axis positioning stage (Thorlabs XRR1, XR25P-K2, PY003), with motorized *x*, *y*, and *z*translation (Thorlabs KDC101 and Z925B) for PSF calibration.

Fluorescence is simultaneously imaged by a reference epi-fluorescence microscope (Nikon Plan Apo *λD* 4×/0.20 NA with a Thorlabs TTL200-A tube lens) positioned beneath the sample and by Bio-CM^2^ positioned above the sample. Both imaging systems are mounted on five-axis alignment stages (Thorlabs PY005). A filtered and expanded 532 nm laser (Thorlabs CPS532) is used for optical alignment.

#### FOV characterization

The designed FOV was experimentally validated using a fluorescent resolution target masked to different effective imaging areas. As predicted by our geometric analysis, the four imaging channels remain completely separated on the sensor within the designed 7.5 × 10 mm^2^ FOV. When the object size exceeds the design limit, adjacent sub-images begin to overlap, in agreement with the derived geometric design constraints (Supplementary Section 1.3).

#### PSF calibration

PSFs were acquired by scanning a 4 µm fluorescent bead (Thermo Fisher Scientific) across the 7.5 mm × 10.0 mm imaging FOV using a motorized translation stage. The bead was sampled on a 31 × 41 grid with a 250 µm step size. Owing to the partially overlapping imaging fields of the four lenses, each lens recorded PSFs at 17 × 23 scan locations, including 14 × 18 locations unique to that lens, 14 × 5 + 3 × 18 locations shared with one adjacent lens, and 3 × 5 locations shared among all four lenses (Supplementary Section 1.4). This procedure yielded a spatially varying PSF dataset spanning the entire imaging FOV for subsequent calibration and image reconstruction.

#### Algorithm implementation

A modular computational pipeline was developed to reconstruct seamless wide-FOV images from the raw Bio-CM^2^ measurements. The reconstruction consists of six sequential modules: denoising, flat-field correction, background removal, geometric distortion correction, deconvolution, and image stitching. Each imaging channel is reconstructed independently through the first five modules before the corrected views are combined into a single wide-FOV image during the final stitching step. The modular design enables individual components to be readily replaced or upgraded without affecting the remaining pipeline. Geometric distortion correction, deconvolution, and stitching rely on the experimentally calibrated PSFs described above. Throughout this section, ***ρ*** denotes the image-space coordinate and ***x*** denotes the corresponding object-space coordinate, unless otherwise specified.

#### Denoising

Starting from the raw measurement *I*_raw_(***ρ***), outliers arising from hot pixels are first removed. For static samples, we apply BM3D [42], which exploits spatial redundancy and transform-domain sparsity for single-image denoising.

For dynamic *in vivo* neural recordings, where neuronal signals are weak and temporally correlated, we instead employ the self-supervised spatiotemporal denoising framework SUPPORT [30]. SUPPORT combines a blind-spot network for extracting spatial features with a U-Net that incorporates temporal information from neighboring frames. The network parameters *θ* are optimized according to

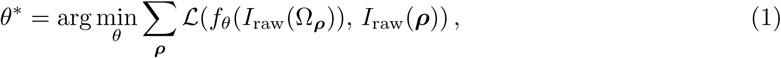

where *f_θ_* denotes the neural network parameterized by *θ*, *L*(·, ·) is the arithmetic mean of the L1 and L2 losses, and Ω***_ρ_*** denotes the spatiotemporal neighborhood centered at ***ρ***. Additional implementation details of SUPPORT are provided in Supplementary Section 2.1. The denoised image is denoted by ^ɪ^_denoised_(***ρ***).

#### Flat-field correction

To compensate for spatially varying illumination and vignetting, a flat-field calibration image *I*_fc_(***ρ***) is acquired by imaging a uniform fluorescent sample (Rhodamine 123 dissolved in water; Thermo Fisher Scientific). The calibration image is normalized to have a unit mean intensity. The denoised image is then corrected according to

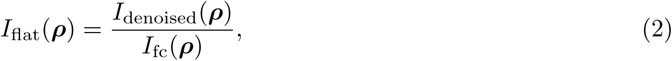

where *I*_flat_(***ρ***) denotes the flat-field-corrected image.

#### Background removal

For static samples, out-of-focus background is suppressed using a modified dark-sectioning algorithm [43]. The algorithm separately estimates the in-focus high-frequency and low-frequency image components before recombining them.

The high-frequency component is inherently in focus and is directly extracted from the flat-field-corrected image using a Gaussian high-pass filter:

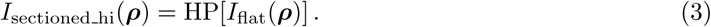

The low-frequency component, however, contains both in-focus structures and out-of-focus background. Applying a complementary Gaussian low-pass filter yields

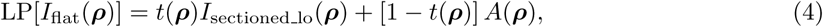

where *A*(***ρ***) denotes the coarse background estimated by further blurring the low-pass image, and *t*(***ρ***) is the spatially varying mixing coefficient.

The coefficient *t*(***ρ***) is estimated using the dark-channel prior. Specifically, because the dark channel computed with a window size matching the width of LP[PSF(***ρ***)] is approximately zero for *I*_sectioned lo_(***ρ***), applying the dark-channel operator to both sides of the equation above enables *t*(***ρ***) to be estimated. The recovered mixing coefficient is then substituted back to obtain *I*_sectioned lo_(***ρ***).

Finally, the sectioned image is synthesized as

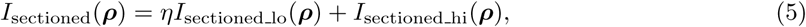

where

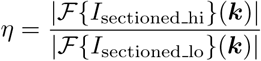

is chosen as the ratio of the Fourier magnitudes at the cutoff frequency ***k*** (by default, half of the system cutoff frequency), providing a smooth spectral transition between the low- and high-frequency components.

#### PSF processing

After PSF calibration, a 180 × 180 pixel patch is extracted around each calibration point. The nominal image-space center of the *m*-th PSF for the *i*-th lens is predicted from the paraxial imaging geometry as

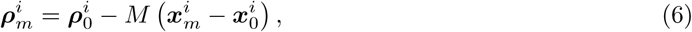

Where 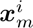 and 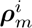 denote the object-space scan position and the corresponding nominal image-space position of the *m*-th calibration point for the *i*-th lens (*i* ∈ {1, 2, 3, 4}), *m* = 0 corresponds to the on-axis point, and *M* is the paraxial magnification.

The nominal position 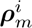 coincides with the PSF centroid only for the on-axis calibration point. For off-axis points, optical distortion shifts the measured PSF centroid to a distorted position, denoted by 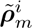. To compensate for this distortion, each off-axis PSF is registered to its nominal position using cross-correlation with the corresponding on-axis PSF as the reference. The registered PSF is denoted by 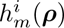.

#### Distortion correction

Using the corresponding undistorted and distorted PSF positions, 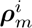 and 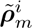, respectively, we parameterize the geometric distortion of each lens with a radial distortion model. For the *i*-th lens, the distorted position predicted from the undistorted coordinate is

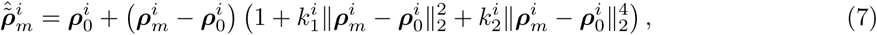

where 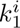 and 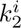 denote the first- and second-order radial distortion coefficients. The coefficients are estimated by minimizing the discrepancy between the predicted and measured distorted PSF positions over all calibration points,

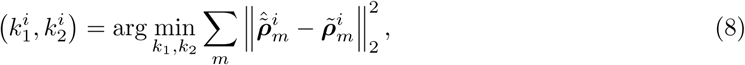

with 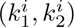 initialized to (0, 0). The fitted distortion model is then inverted to map each pixel in the distorted sub-image 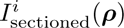 onto the undistorted coordinate system by bilinear interpolation, yielding the corrected sub-image 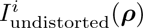

#### Patch-wise deconvolution

To account for the spatially varying PSF while maintaining computational efficiency, we partition 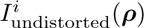 into overlapping patches 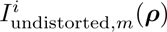 centered at 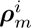 . Each patch is deconvolved using Tikhonov regularization,

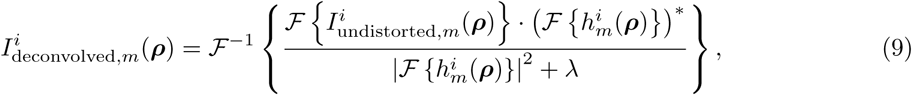

where ℱ{·} and ℱ^−1^{·} denote the Fourier and inverse Fourier transforms, respectively, (·)^∗^ denotes complex conjugation, and *λ* is the Tikhonov regularization parameter. The deconvolved patches are finally merged using alpha blending to suppress boundary artifacts, yielding the deconvolved sub-image 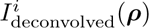

#### Stitching

For quasi-two-dimensional samples, such as the resolution target and fixed brain slices, the four deconvolved sub-images are directly merged using alpha blending to generate the final reconstruction, *I*_stitched_(***ρ***).

For specimens with appreciable depth variation, such as the curved mouse cortex, parallax introduces local misalignment between overlapping views. To compensate for this effect, corresponding feature points are manually selected between neighboring sub-images, from which pixel-wise warp fields are generated by linear interpolation. The resulting warp fields are applied to 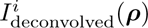 prior to alpha blending, enabling seamless stitching of the four sub-images (Supplementary Section 2.2).

### Sample preparation

#### Fixed brain slice

This study is performed in strict accordance with the recommendations in the Guide for the Care and Use of Laboratory Animals of the National Institutes of Health. All animals are handled according to approved Institutional Animal Care and Use Committee (IACUC) protocols (#201800540) of Boston University.

Ex vivo fixed rodent brain slices with GFP-expressing neurons in the bed nucleus of the stria terminalis (BNST) are prepared using viral-mediated labeling in male C57BL/6 mice. Following anesthesia and preoperative analgesia, a small bilateral craniotomy is performed over each injection site. Viral solutions (100–200 nL; pAAV-CAG-GFP, Addgene #37825-AAVrg) are delivered to the BNST, labeling local neurons at the injection site as well as upstream brain areas. After recovery, the animals are anesthetized, euthanized, and perfused with paraformaldehyde, and their brains are sectioned coronally at 75 µm thickness. The sections are then mounted for imaging.

##### Caenorhabditis elegans

Transgenic nematodes AM141 [unc-54p::Q40::YFP], expressing a 40-polyglutamine repeat fused to yellow fluorescent protein (YFP), are cultured on nematode growth medium (NGM) plates seeded with OP50 *Escherichia coli*. The NGM contains 20 mM phosphate buffer, 0.8 mM CaCl_2_, 0.8 mM MgSO_4_, 17.5 g/L agar, 3.0 g/L NaCl, 2.5 g/L peptone, and 0.005 g/L cholesterol. Worms are maintained at 15 ^◦^C. Young adult worms (day 1–3 adults) are placed on a flat ∼20 µL 2% agar pad for imaging.

#### Mice

All animal experiments are conducted in compliance with the National Institutes of Health Guide for the Care and Use of Laboratory Animals and are approved by the Boston University Institutional Animal Care and Use Committee. Mice are housed under a 12-h light/dark cycle with lights on at 7:30 a.m., with standard rodent chow and water provided *ad libitum*.

For cortical neuronal imaging, we use adult (6–12-week-old) VIP-Cre (The Jackson Laboratory, strain #010908); Ai162 (strain #031562) mice, which express GCaMP6s in VIP-expressing interneurons. For vascular imaging, fluorescein isothiocyanate (FITC)-dextran (2 MDa, MilliporeSigma) was injected intravenously under isoflurane anesthesia 15–30 min before imaging to label the cerebral vasculature. The full-cortex cranial window surgery followed a previously described procedure [24]. Briefly, dexamethasone (4.8 mg kg^−1^), extended-release buprenorphine (EthiqaXR, 3.25 mg kg^−1^), and meloxicam (5 mg kg^−1^) are administered approximately 4 h before surgery. Mice are anesthetized with either isoflurane (2% in O_2_ for induction and 1% in O_2_ throughout the procedure) or ketamine/xylazine, and body temperature is maintained at 37^◦^C throughout surgery. A bilateral chronic glass cranial window over the dorsal cortex was created by replacing the dorsal cranium with two pieces of flat glass, one over each hemisphere, and securing a custom-designed titanium headpost to the cranium. Mice recover for at least one week after surgery before habituation to head fixation, and for at least three weeks before imaging experiments begin. Both vascular and neuronal imaging were performed in the same animal through the same chronic bilateral glass cranial window.

For olfactory bulb imaging, mice are obtained from the Jackson Laboratory (Tbet-Cre driver and Ai148 conditional reporter; stock #s 024507 and 030328, respectively) and crossed to achieve selective expression of GCaMP6f in second-order neurons of the main olfactory bulb. Mice are group-housed prior to implantation and then singly housed afterward to prevent damage to the cranial window. For surgery, animals are anesthetized with isoflurane (∼3% for induction, 1.5% for maintenance, delivered in pure O_2_ at ∼1.5 L/min) and placed in a stereotaxic frame. The skin overlying the skull is shaved and treated with topical lidocaine (0.5%), followed by iodine and 70% isopropyl alcohol. The skin above the olfactory bulb and anterior cortex is carefully removed, and an intact-skull window is constructed by thinning the bone over the dorsal olfactory bulbs to a thickness of ∼ 100 µm. The thinned bone is covered with a thin layer of cyanoacrylate (Vetbond) to reduce scattering and improve optical access, followed by optical adhesive (Norland type 81) and a 4 mm glass coverslip (Warner Instruments). A stainless steel headpost is attached immediately posterior to the cranial window for later head fixation, and both the headpost and window are attached with dental acrylic (Metabond).

### Experimental settings

#### *In vivo* head-fixed mouse imaging with Bio-CM**^2^**

During imaging, Bio-CM^2^ was mounted on a compact 5-axis translation stage (Thorlabs PY005) to enable fine adjustment of the imaging position. The head-fixed mouse was placed on a custom-built platform mounted on motorized translation stages (Thorlabs MLJ150), allowing independent alignment of the animal with respect to the imaging system. Because Bio-CM^2^ provides a large DOF (400–500 µm), the imaging plane could be selected by a single axial positioning step prior to recording, without the need for axial scanning during data acquisition. Once the desired focal plane was established, videos were acquired continuously at 10 Hz. For neuronal calcium imaging, recordings lasted 3 min. For vascular imaging, recordings lasted 5 min after intravenous FITC-dextran injection, which served as the primary fluorescence contrast for visualizing the cerebral vasculature.

For MOB imaging, video is recorded at 20 Hz for ∼ 30 s. We begin odor stimulation ∼ 5 s after the start of the recording. Odors are presented manually for 4 s using 50 µL of odorant solution applied to a sterile cotton swab positioned 1 cm in front of the animal’s nose (either pentyl acetate or benzaldehyde, diluted in mineral oil to final concentrations of 2% and 10%, respectively).

#### Two-photon *in vivo* head-fixed mouse imaging

Two-photon imaging was performed in awake, head-fixed mice using a commercial laser-scanning microscope (Bruker Ultima Investigator Plus) equipped with a 20×/1.0 NA objective (Olympus XLUM- PlanFLNXW) and a Ti:Sapphire excitation laser (Coherent Chameleon Discovery, *λ* = 920 nm). Image stacks were acquired with a 5 µm axial step size over a depth range of 0–350 µm from the cortical surface.

To estimate the effective imaging depth of Bio-CM^2^ under *in vivo* scattering conditions, maximum-intensity projections (MIPs) were generated from different depth ranges of the two-photon image stack and compared with the corresponding Bio-CM^2^ reconstruction. The highest structural correspondence was obtained using a 100–300 µm depth projection, which was therefore used for comparison in Supplementary Section 5. This result indicates that, despite operating as a one-photon imaging modality, Bio-CM^2^ is able to capture neuronal activity covering most of the cortical layer 2/3 under sparse labeling conditions.

### Data analysis

#### Vessel diameter analysis

For quantitative vasomotion analysis, arterial vessel segments were manually selected from the reconstructed images based on their morphology and branching pattern. Venous vessels were excluded because their diameter fluctuations primarily reflect passive hemodynamic changes rather than active vasomotion.

Vessel diameter was estimated from the fluorescence intensity profile measured along a manually selected line perpendicular to the vessel axis. The vessel boundaries were identified as the locations of the maximum positive and negative intensity gradients, and the vessel diameter was defined as the distance between these two boundary points. Repeating this procedure for each frame yielded a time-varying diameter trace, which was normalized to its mean value to obtain the relative diameter change (Δ*d/d*). The resulting traces were used to quantify vasomotion and compute the corresponding power spectra.

#### Single-neuron calcium signal extraction

Calcium neuronal signals were extracted from the reconstructed full-FOV videos using the CNMF-E algorithm [31, 32] (Supplementary Section 4.1). CNMF-E models the recorded fluorescence movie as the sum of neuronal signals, background fluorescence, and residual noise. Each neuronal component is represented by a spatial footprint and its corresponding temporal fluorescence trace, while the background is decomposed into a constant baseline and temporally varying fluctuations.

After CNMF-E fitting, the detected components were subjected to quality control. Candidate components located outside manually annotated cortical regions were discarded. The remaining components were further filtered using the spatial correlation metric provided by CNMF-E, which quantifies the agreement between the inferred spatial footprint and the corresponding fluorescence pattern in the raw imaging data. Only components satisfying both criteria were retained for subsequent analysis.

The CNMF-E parameter selection and neuron identification were independently cross-validated using a simple local-maximum intensity detection method (Supplementary Section 4.2).

The temporal fluorescence traces of the retained neurons were used for all downstream analyses. Calcium-response kinetics were quantified by measuring the time to peak (*t*_peak_) and the post-peak half-decay time (*t*_1*/*2_) for individual calcium transients.

#### Regional hemodynamics and neuronal activity

To quantify mesoscale cortical dynamics, a dorsal projection of the Allen Mouse Brain Atlas [44] was registered to the reconstructed FOV using prominent vascular and cortical landmarks. Twenty-eight anatomical parcels spanning the bilateral dorsal cortex were defined according to the registered atlas and numbered as follows: 1, RSPd-R; 2, RSPagl-R; 3, VISp-R; 4, VISa-R; 5, VISam-R; 6, VISrl-R; 7, VISpm-R; 8, SSp-un-R; 9, SSp-ul-R; 10, SSp-ll-R; 11, SSp-tr-R; 12, SSp-bfd-R; 13, MOs-R; 14, MOp-R; 15, RSPd-L; 16, RSPagl-L; 17, VISp-L; 18, VISa-L; 19, VISam-L; 20, VISrl-L; 21, VISpm-L; 22, SSp-un-L; 23, SSp-ul-L; 24, SSp-ll-L; 25, SSp-tr-L; 26, SSp-bfd-L; 27, MOs-L; and 28, MOp-L, where R and L denote the right and left hemispheres, respectively.

For each cortical parcel, the fluorescence signal was computed by spatially averaging pixels within the anatomical parcel that overlapped the manually delineated cortical region imaged by Bio-CM^2^. The resulting regional time series were converted to relative fluorescence changes (Δ*F/F* ) by normalizing to their mean fluorescence intensity. These regional activity traces were subsequently used to quantify cortex-wide spatiotemporal dynamics through pairwise correlation analysis and seed-based functional connectivity mapping [45].

#### Glomerular activity analysis

The reconstructed video was converted to relative fluorescence changes (Δ*F/F* ) by normalizing each frame to a baseline image computed as the average of the first ∼5 s of recording. An MIP of the Δ*F/F* video was generated for each odor stimulus, and regions of interest (ROIs) corresponding to individual glomeruli were manually selected from the MIP images. For each ROI, the glomerular activity trace was computed as the mean Δ*F/F* value of all pixels within the ROI at each time point.

## Acknowledgment

We acknowledge the support by the Neurophotonics Center at Boston University. We thank Qianwan Yang, Tongyu Li, Shuying Li, and Yujia Xue for helpful discussions on Bio-CM^2^. We thank Sangsoo Lee for assistance with extracting additional vessel ROIs. We thank the Boston University Shared Computing Cluster for providing computational resources.

---

**Extended Data Figure 1.**
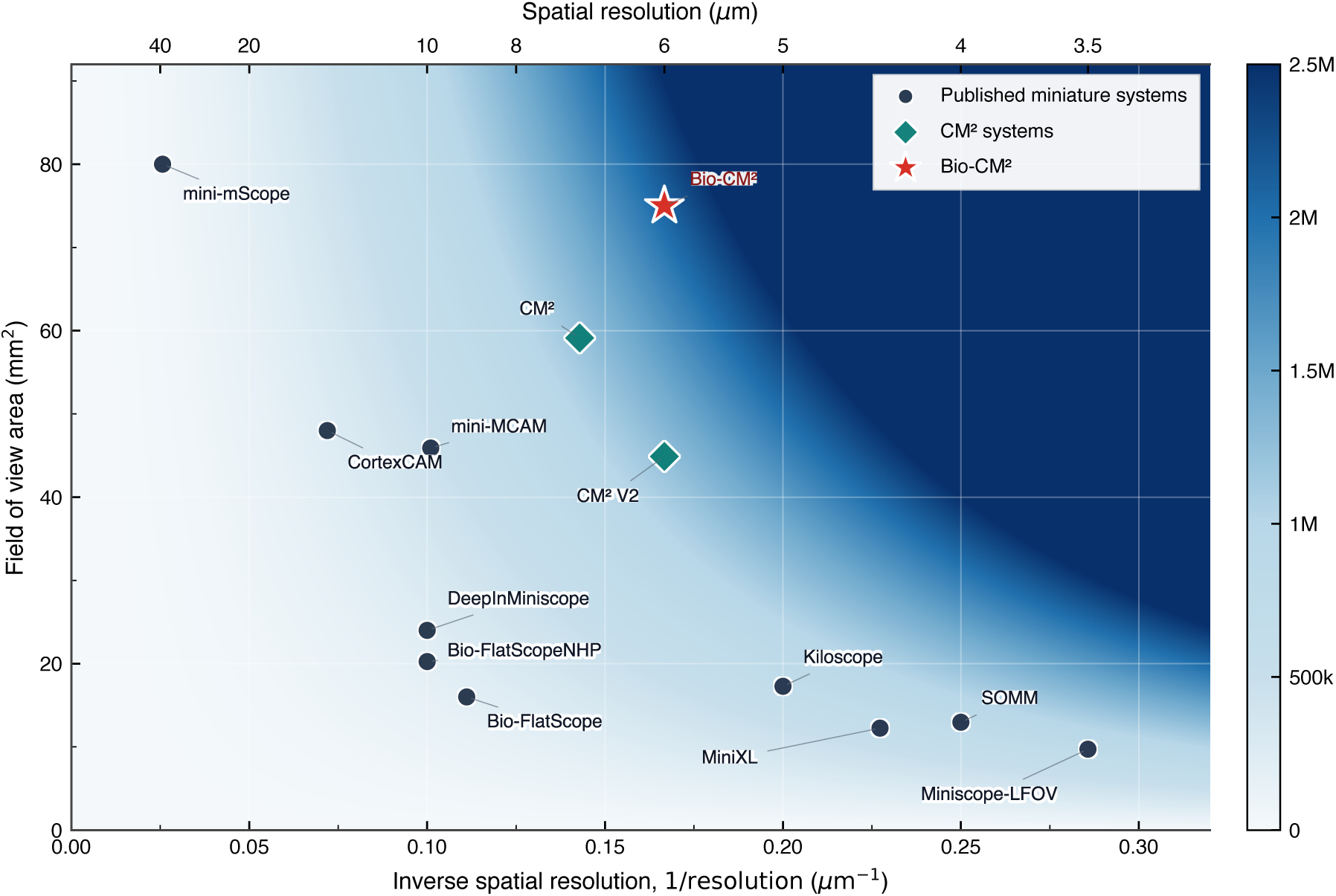
FOV and best achievable spatial resolution of miniature imaging systems. FOV area is plotted against lateral resolution for representative published miniature imaging systems and Bio-CM^2^. Because the spatial resolution of miniature imaging systems generally varies across the imaging FOV (Supplementary Section 1.8), each point represents the best lateral resolution reported in the corresponding work. The background contours indicate constant values of a FOV–resolution figure of merit, defined as the FOV area divided by the square of the lateral resolution. Bio-CM^2^ combines a large FOV with high spatial resolution, placing it near the upper-left region of the comparison space.

**Extended Data Figure 2.**
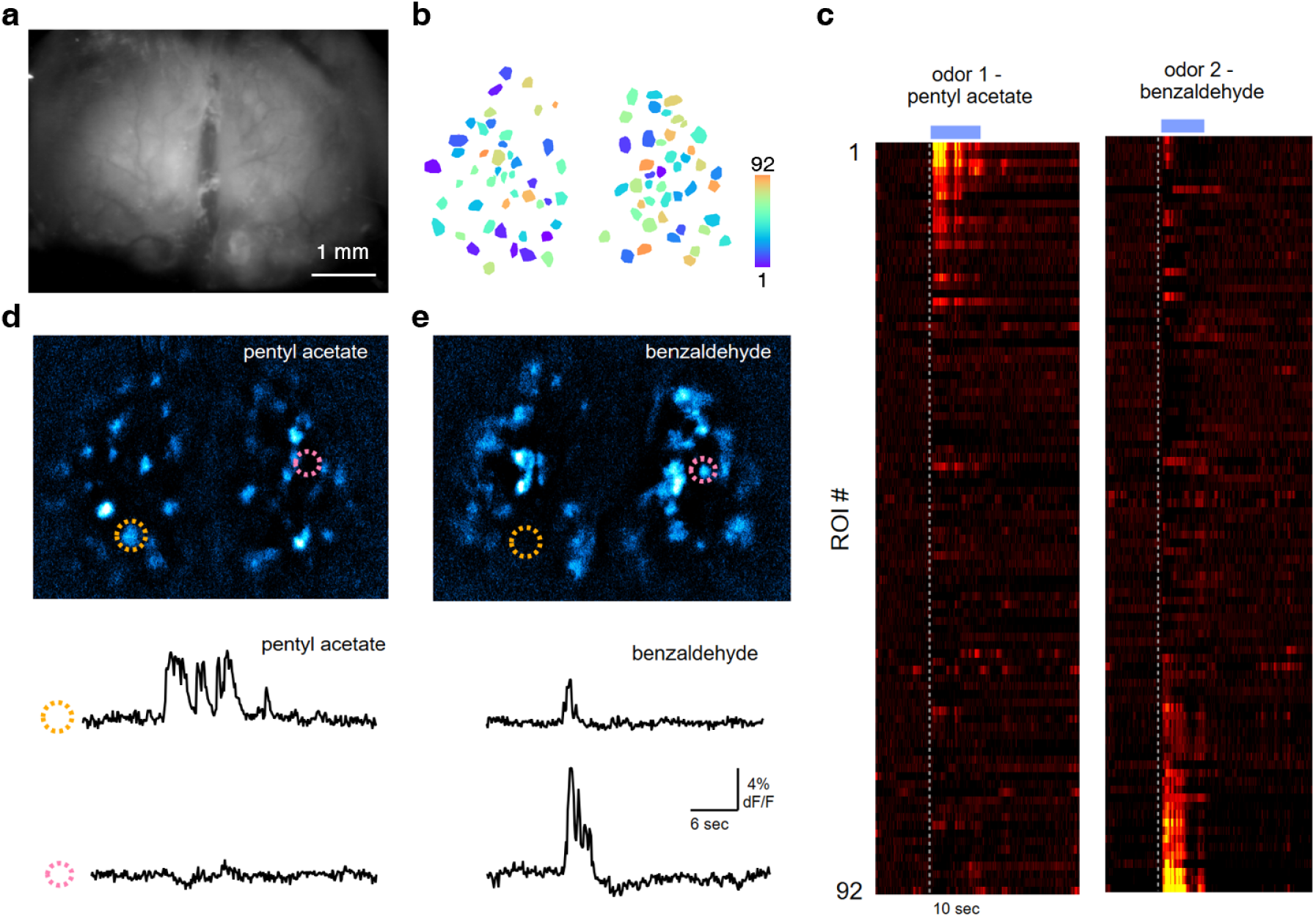
Odor-evoked glomerular activity in the mouse MOB. (a) Reconstructed fluorescence image of the dorsal surface of the MOB acquired with Bio-CM^2^. (b) 92 manually selected glomerular ROIs used for activity analysis. Colors indicate ROI indices. (c) Δ*F/F* responses of all glomerular ROIs to two odorants, pentyl acetate and benzaldehyde. Each row corresponds to one ROI. The dashed line indicates odor onset, and the blue bar denotes the odor presentation period. (d,e,) Odor-evoked Δ*F/F* activity maps for pentyl acetate (d) and benzaldehyde (e), illustrating distinct glomerular activation patterns. Representative glomeruli are marked by colored circles, and their corresponding Δ*F/F* traces are shown below.

**Extended Data Figure 3.**
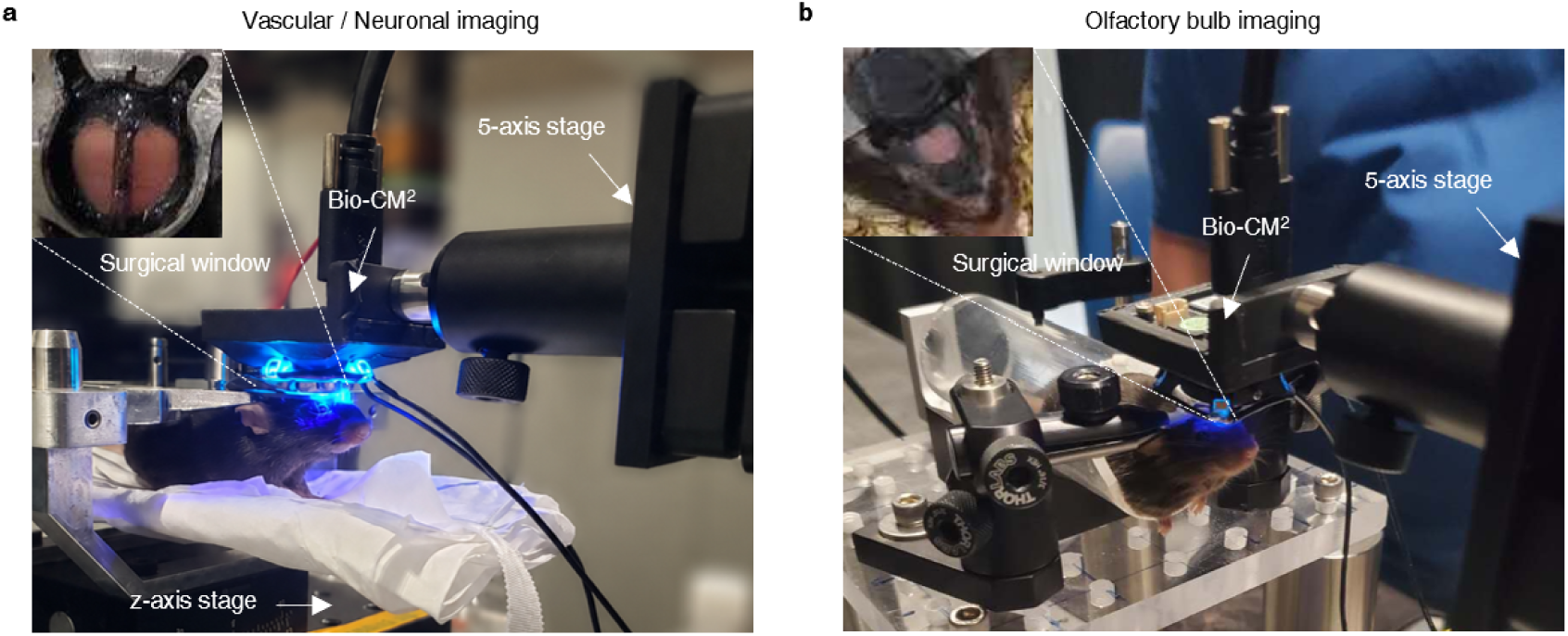
Experimental setups for *in vivo* mouse imaging. (a) Experimental setup for cortex-wide vascular and neuronal imaging through a chronic cranial window. Inset, top view of the chronic cranial window spanning the entire dorsal cortex. (b) Experimental setup for calcium imaging of the main olfactory bulb (MOB). Inset, top view of the exposed dorsal surface of the bilateral olfactory bulbs.

## References

[1] Nicholas A. Steinmetz, Peter Zatka-Haas, Matteo Carandini, and Kenneth D. Harris. Distributed coding of choice, action and engagement across the mouse brain. Nature, 576(7786):266–273, 2019.

[2] Jingtao Fan, Jinli Suo, Jiamin Wu, Hao Xie, Yibing Shen, Feng Chen, Guijin Wang, Liangcai Cao, Guofan Jin, Quansheng He, et al. Video-rate imaging of biological dynamics at centimetre scale and micrometre resolution. Nature Photonics, 13(11):809–816, 2019.

[3] Isaac V. Kauvar, Timothy A. Machado, Elle Yuen, John Kochalka, Minseung Choi, William E. Allen, Gordon Wetzstein, and Karl Deisseroth. Cortical observation by synchronous multifocal optical sampling reveals widespread population encoding of actions. Neuron, 107(2):351–367.e19, 2020.

[4] Mark Harfouche, Kanghyun Kim, Kevin C. Zhou, Pavan Chandra Konda, Sunanda Sharma, Eric E. Thomson, Colin Cooke, Shiqi Xu, Lucas Kreiss, Amey Chaware, Xi Yang, Xing Yao, Vinayak Pathak, Martin Bohlen, Ron Appel, Aur’elien B‘egue, Clare Cook, Jed Doman, John Efromson, Gregor Horstmeyer, Jaehee Park, Paul Reamey, Veton Saliu, Eva Naumann, and Roarke Horstmeyer. Imaging across multiple spatial scales with the multi-camera array microscope. Optica, 10(4):471–480, 2023.

[5] Yuanlong Zhang, Mingrui Wang, Qiyu Zhu, Yuduo Guo, Bo Liu, Jiamin Li, Xiao Yao, Chui Kong, Yi Zhang, Yuchao Huang, Hai Qi, Jiamin Wu, Zengcai V. Guo, and Qionghai Dai. Long-term mesoscale imaging of 3d intercellular dynamics across a mammalian organ. Cell, 187(21):6104–6122.e25, 2024.

[6] Nicholas James Sofroniew, Daniel Flickinger, Jonathan King, and Karel Svoboda. A large field of view two-photon mesoscope with subcellular resolution for in vivo imaging. eLife, 5:e14472, 2016.

[7] Daniel Aharoni, Baljit S Khakh, Alcino J Silva, and Peyman Golshani. All the light that we can see: a new era in miniaturized microscopy. Nature methods, 16(1):11–13, 2019.

[8] Adolf W. Lohmann. Scaling laws for lens systems. Applied Optics, 28(23):4996–4998, 1989.

[9] Changliang Guo, Garrett J Blair, Megha Sehgal, Federico N Sangiuliano Jimka, Arash Bellafard, Alcino J Silva, Peyman Golshani, Michele A Basso, Hugh Tad Blair, and Daniel Aharoni. Miniscope-lfov: A large-field-of-view, single-cell-resolution, miniature microscope for wired and wire-free imaging of neural dynamics in freely behaving animals. Science advances, 9(16):eadg3918, 2023.

[10] Pingping Zhao, Changliang Guo, Mian Xie, Liangyi Chen, Peyman Golshani, and Daniel Aharoni. MiniXL: An open-source, large field-of-view epifluorescence miniscope enabling single-cell resolution and multi-region imaging in mice. Science Advances, 11(24):ads4995, 2025.

[11] Yuanlong Zhang, Lekang Yuan, Qiyu Zhu, Jiamin Wu, Tobias N”obauer, Rujin Zhang, Guihua Xiao, Mingrui Wang, Hao Xie, Zengcai Guo, Qionghai Dai, and Alipasha Vaziri. A miniaturized mesoscope for the large-scale single-neuron-resolved imaging of neuronal activity in freely behaving mice. Nature Biomedical Engineering, 8(6):754–774, 2024.

[12] Josefa R. Scherrer, Galen F. Lynch, Jie J. Zhang, and Michale S. Fee. An optical design enabling lightweight and large field-of-view head-mounted microscopes. Nature Methods, 20(4):546–549, 2023.

[13] Jia Hu, Arun Cherkkil, Daniel A Surinach, Ibrahim Oladepo, Ridwan Hossain, Skylar Fausner, Kapil Saxena, Eunsong Ko, Ryan Peters, Michael Feldkamp, et al. Pan-cortical cellular imaging in freely behaving mice using a miniaturized micro-camera array microscope (mini-mcam). Science Advances, 11(44):eadt3634, 2025.

[14] Arun Cherkkil, Zoey Viavattine, Vamsy Kota, Kapil Saxena, Daniel Surinach, Ibrahim Oladepo, Jill Juneau, Jia Hu, Skylar M.L. Fausner, Eunsong Ko, Shubham Panchal, Sanjana Srivatsa, Malachi R. Lehman, Roarke Horstmeyer, and Suhasa B Kodandaramaiah. Cortex-wide cellular imaging in freely locomoting mice using cortex camera array microscope (cortexcam). bioRxiv, 2026.

[15] Jesse K Adams, Dong Yan, Jimin Wu, Vivek Boominathan, Sibo Gao, Alex V Rodriguez, Soonyoung Kim, Jennifer Carns, Rebecca Richards-Kortum, Caleb Kemere, et al. In vivo lensless microscopy via a phase mask generating diffraction patterns with high-contrast contours. Nature Biomedical Engineering, 6(5):617–628, 2022.

[16] Jimin Wu, Yuzhi Chen, Ashok Veeraraghavan, Eyal Seidemann, and Jacob T Robinson. Mesoscopic calcium imaging in a head-unrestrained male non-human primate using a lensless microscope. Nature Communications, 15(1):1271, 2024.

[17] Kyrollos Yanny, Nick Antipa, William Liberti, Sam Dehaeck, Kristina Monakhova, Fanglin Linda Liu, Konlin Shen, Ren Ng, and Laura Waller. Miniscope3D: optimized single-shot miniature 3d fluorescence microscopy. Light: Science Applications, 9(1):171, 2020.

[18] Robert Prevedel, Young-Gyu Yoon, Maximilian Hoffmann, Nikita Pak, Gordon Wetzstein, Saul Kato, Tina Schr”odel, Ramesh Raskar, Manuel Zimmer, Edward S. Boyden, and Alipasha Vaziri. Simultaneous whole-animal 3d imaging of neuronal activity using light-field microscopy. Nature Methods, 11(7):727–730, 2014.

[19] Yujia Xue, Ian G Davison, David A Boas, and Lei Tian. Single-shot 3d wide-field fluorescence imaging with a computational miniature mesoscope. Science advances, 6(43):eabb7508, 2020.

[20] Yujia Xue, Qianwan Yang, Guorong Hu, Kehan Guo, and Lei Tian. Deep-learning-augmented computational miniature mesoscope. Optica, 9(9):1009–1021, 2022.

[21] Qianwan Yang, Ruipeng Guo, Guorong Hu, Yujia Xue, Yunzhe Li, and Lei Tian. Wide-field, high-resolution reconstruction in computational multi-aperture miniscope using a fourier neural network. Optica, 11(6):860–871, 2024.

[22] Feng Tian, Ben Mattison, and Weijian Yang. Deepinminiscope: Deep learning–powered physics-informed integrated miniscope. Science Advances, 11(37):eadr6687, 2025.

[23] Mathew L Rynes, Daniel A Surinach, Samantha Linn, Michael Laroque, Vijay Rajendran, Judith Dominguez, Orestes Hadjistamoulou, Zahra S Navabi, Leila Ghanbari, Gregory W Johnson, Mojtaba Nazari, Majid H Mohajerani, and Suhasa B Kodandaramaiah. Miniaturized head-mounted microscope for whole-cortex mesoscale imaging in freely behaving mice. Nature Methods, 18(4):417–425, 2021.

[24] Bradley C Rauscher, Natalie Fomin-Thunemann, Sreekanth Kura, Patrick R Doran, Pablo D Perez, Kıvılcım Kılıtç, Emily A Martin, Dora Balog, Nathan X Chai, Francesca A Froio, et al. The neurovascular impulse response function differentially reflects intrinsic neuromodulation across cortical regions. Nature neuroscience, pages 1–9, 2026.

[25] Anna Devor, Peifang Tian, Nozomi Nishimura, Ivan C Teng, Elizabeth MC Hillman, SN Narayanan, Istvan Ulbert, David A Boas, David Kleinfeld, and Anders M Dale. Suppressed neuronal activity and concurrent arteriolar vasoconstriction may explain negative blood oxygenation level-dependent signal. Journal of Neuroscience, 27(16):4452–4459, 2007.

[26] Celine Mateo, Per M Knutsen, Philbert S Tsai, Andy Y Shih, and David Kleinfeld. Entrainment of arteriole vasomotor fluctuations by neural activity is a basis of blood-oxygenation-level-dependent “resting-state” connectivity. Neuron, 96(4):936–948, 2017.

[27] Thomas Broggini, Jacob Duckworth, Xiang Ji, Rui Liu, Xinyue Xia, Philipp Mächler, Iftach Shaked, Leon Paul Munting, Satish Iyengar, Michael Kotlikoff, et al. Long-wavelength traveling waves of vasomotion modulate the perfusion of cortex. Neuron, 112(14):2349–2367, 2024.

[28] Peifang Tian, Anna Devor, Sava Sakadžić, Anders M Dale, and David A Boas. Monte carlo simulation of the spatial resolution and depth sensitivity of two-dimensional optical imaging of the brain. Journal of biomedical optics, 16(1):016006–016006, 2011.

[29] S Shahsavarani, DN Thibodeaux, W Xu, SH Kim, F Lodgher, C Nwokeabia, M Cambareri, AJ Yagielski, HT Zhao, DA Handwerker, et al. Cortex-wide neural dynamics predict behavioral states and provide a neural basis for resting-state dynamic functional connectivity. cell rep. 42, 112527, 2023.

[30] Minho Eom, Seungjae Han, Pojeong Park, Gyuri Kim, Eun-Seo Cho, Jueun Sim, Kang-Han Lee, Seonghoon Kim, He Tian, Urs L Böhm, et al. Statistically unbiased prediction enables accurate denoising of voltage imaging data. Nature Methods, 20(10):1581–1592, 2023.

[31] Pengcheng Zhou, Shanna L Resendez, Jose Rodriguez-Romaguera, Jessica C Jimenez, Shay Q Neufeld, Andrea Giovannucci, Johannes Friedrich, Eftychios A Pnevmatikakis, Garret D Stuber, Rene Hen, et al. Efficient and accurate extraction of in vivo calcium signals from microendoscopic video data. elife, 7:e28728, 2018.

[32] Andrea Giovannucci, Johannes Friedrich, Pat Gunn, Jéŕemie Kalfon, Brandon L Brown, Sue Ann Koay, Jiannis Taxidis, Farzaneh Najafi, Jeffrey L Gauthier, Pengcheng Zhou, et al. Caiman an open source tool for scalable calcium imaging data analysis. elife, 8:e38173, 2019.

[33] Tsai-Wen Chen, Trevor J Wardill, Yi Sun, Stefan R Pulver, Sabine L Renninger, Amy Baohan, Eric R Schreiter, Rex A Kerr, Michael B Orger, Vivek Jayaraman, et al. Ultrasensitive fluorescent proteins for imaging neuronal activity. Nature, 499(7458):295–300, 2013.

[34] Elizabeth M. C. Hillman. Coupling mechanism and significance of the BOLD signal: a status report. Annual Review of Neuroscience, 37:161–181, 2014.

[35] Bruno Cauli, Xin-Kang Tong, Armelle Rancillac, Nella Serluca, Bertrand Lambolez, Jean Rossier, and Edith Hamel. Cortical GABA interneurons in neurovascular coupling: relays for subcortical vasoactive pathways. Journal of Neuroscience, 24(41):8940–8949, 2004.

[36] Ying Ma, Mohammed A. Shaik, Mariel G. Kozberg, Seong-Gi Kim, Jo ao P. Portes, David Timerman, and Elizabeth M. C. Hillman. Resting-state hemodynamics are spatiotemporally coupled to synchronized and symmetric neural activity in excitatory neurons. Proceedings of the National Academy of Sciences, 113(52):E8463–E8471, 2016.

[37] Naoshige Uchida, Yuji K. Takahashi, Manabu Tanifuji, and Kensaku Mori. Odor maps in the mammalian olfactory bulb: domain organization and odorant structural features. Nature Neuroscience, 3(10):1035–1043, 2000.

[38] Zhe Dong, Yu Feng, Keziah Diego, Austin M. Baggetta, Brian M. Sweis, Zachary T. Pennington, Sophia I. Lamsifer, Yosif Zaki, Federico Sangiuliano, Paul A. Philipsberg, Denisse Morales-Rodriguez, Daniel Kircher, Paul Slesinger, Tristan Shuman, Daniel Aharoni, and Denise J. Cai. Simultaneous two-color imaging with a dual-channel miniscope in freely behaving mice. Science Advances, 11(27):eadr6470, 2025.

[39] Patrick R Doran, Natalie Fomin-Thunemann, Rockwell P Tang, Dora Balog, Bernhard Zimmerman, Kıvılcım Kılıç, Emily A Martin, Sreekanth Kura, Harrison P Fisher, Grace Chabbott, et al. Widefield in vivo imaging system with two fluorescence and two reflectance channels, a single scmos detector, and shielded illumination. Neurophotonics, 11(3):034310–034310, 2024.

[40] Qianwan Yang, Zhixiong Chen, Jiaqi Zhang, Ruipeng Guo, Guorong Hu, and Lei Tian. Coordinate-conditioned deconvolution for scalable spatially varying high-throughput imaging. *arXiv preprint arXiv:2602.01065*, 2026.

[41] Yuanlong Zhang, Guoxun Zhang, Xiaofei Han, Jiamin Wu, Ziwei Li, Xinyang Li, Guihua Xiao, Hao Xie, Lu Fang, and Qionghai Dai. Rapid detection of neurons in widefield calcium imaging datasets after training with synthetic data. Nature Methods, 20(5):747–754, 2023.

[42] Kostadin Dabov, Alessandro Foi, Vladimir Katkovnik, and Karen Egiazarian. Image denoising by sparse 3-d transform-domain collaborative filtering. IEEE Transactions on image processing, 16(8):2080–2095, 2007.

[43] Ruijie Cao, Yaning Li, Yao Zhou, Meiqi Li, Fangrui Lin, Wenyi Wang, Guoxun Zhang, Gang Wang, Boya Jin, Wei Ren, et al. Dark-based optical sectioning assists background removal in fluorescence microscopy. Nature Methods, 22(6):1299–1310, 2025.

[44] Quanxin Wang, Song-Lin Ding, Yang Li, Josh Royall, David Feng, Phil Lesnar, Nile Graddis, Maitham Naeemi, Benjamin Facer, Anh Ho, et al. The allen mouse brain common coordinate framework: a 3d reference atlas. Cell, 181(4):936–953, 2020.

[45] Simon Musall, Matthew T. Kaufman, Ashley L. Juavinett, Steven Gluf, and Anne K. Churchland. Single-trial neural dynamics are dominated by richly varied movements. Nature Neuroscience, 22(10):1677–1686, 2019.

